# Whole transcriptome analysis reveals ELK3 as a key driver of metastasis through regulation of 3D migration and stemness in triple-negative breast cancer cells

**DOI:** 10.64898/2026.07.21.739737

**Authors:** Daniel Cruceriu, Loredana Balacescu, Oana Sava, Stefan Miron, Irma-Lidia Szigyarto, Alexandrina Burlacu, Manuela Banciu, Ovidiu Balacescu

**Author notes:** Corresponding author: Daniel Cruceriu, PhD, “Babes-Bolyai” University, Department of Molecular Biology and Biotechnology, 5-7 Clinicilor Street, 400006 Cluj-Napoca, Romania (primary affiliation), The Oncology Institute “Prof. Dr. Ion Chiricuta”, Department of Genetics, Genomics and Experimental Pathology, 34-36 Republicii Street, 400015 Cluj-Napoca, Romania.

## Abstract

Metastasis is the leading cause of mortality in breast cancer and remains largely untargeted therapeutically. Identifying molecular drivers of metastatic progression is essential for developing effective treatments. This study investigated the role of the transcription factor ELK3 in triple-negative breast cancer (TNBC) metastasis by defining the cellular and molecular processes it regulates. MDA231 cells with ELK3 overexpression (OE) or knockdown (KD) were generated by lentiviral transduction. Transcriptomic alterations induced by ELK3-KD were analyzed by microarray and validated by RT-qPCR. Ingenuity Pathway Analysis and Gene Set Enrichment Analysis identified ELK3-dependent metastasis-associated pathways, which were functionally validated using 3D microfluidic migration assays, mammosphere formation assays, and flow cytometry/ AlamarBlue proliferation assays. High ELK3 expression correlated with a mesenchymal phenotype in BC cell lines and lymph node invasion in patient tumors. ELK3-KD significantly altered 740 genes, many linked to migration and stemness. Functionally, ELK3 enhanced 3D confined migration, likely through regulation of EMT, cell adhesion and protrusion formation. ELK3 also promoted cancer stem cell traits, potentially via hypoxia-related and WNT/β-catenin, JAK/STAT3, TGF-β, Notch1, and NF-κB signaling pathways. Additionally, ELK3 induced cellular quiescence while suppressing proliferation under adherent conditions. Overall, ELK3 acts as a pro-metastatic regulator in TNBC by promoting migration and stemness.

## I. Introduction

Breast cancer (BC) is the most commonly diagnosed cancer among women worldwide, accounting for over 2.2 million new cases in 2022 and ranking as the fourth leading cause of cancer-related mortality^1^. As a highly heterogeneous disease, BC is classified into four intrinsic molecular subtypes, luminal A, luminal B, HER2-positive, and triple negative, each with distinct clinical behaviors and treatment responses^2^. Among these, triple negative breast cancer (TNBC), which constitutes approximately 15-25% of all BC cases, is the most aggressive subtype, marked by poor prognosis and a lack of targeted therapies due to the absence of estrogen, progesterone, and HER2 receptors^3^.

Metastasis remains the primary cause of death in BC patients, yet effective molecularly targeted treatments for this process are lacking^4^. TNBC is characterized by the highest metastatic potential among BC molecular subtypes and the lowest five-year survival rate if the tumor has spread into nearby lymph nodes (60-65% survival rate) or distant sites (10-12% survival rate)^5^. Identifying novel molecular drivers of metastasis in TNBC is therefore imperative to advance the development of new therapeutic strategies.

ELK3, also known as Net or Sap2, is a member of the ETS-domain transcription factor family and functions primarily as a transcriptional repressor under basal conditions, although it can act as a transcriptional activator in response to RAS/ERK and p38/MAPK signaling^6^. ELK3 expression is elevated in tumors compared to adjacent healthy tissues across various cancer types, and its upregulation is associated with poorer overall survival^7–9^. In breast cancer, higher ELK3 levels correlate with a more aggressive phenotype, with significant overexpression observed in TNBC relative to luminal subtypes^10^.

ELK3 is involved in several oncogenic processes, including cell motility, proliferation, survival, immune response, and chemotherapy resistance^6^. Particularly relevant to metastasis are processes such as cell migration, invasion, adhesion, and epithelial-to-mesenchymal transition (EMT), all of which are promoted by ELK3 across cancer types^6^. Additionally, in specific cancers such as glioma^11^ and colorectal cancer^12^, ELK3 appears to enhance cancer cell stemness. Cancer stem cells (CSCs), a subpopulation within tumors with self-renewal capacity, drive metastasis and contribute to chemoresistance^13^. In TNBC, ELK3 has been shown to induce a mesenchymal phenotype^14^, repressing E-cadherin expression^15^, and promote cell migration, invasion^10^, adhesion^16^, and extravasation^17^. Furthermore, ELK3 overexpression has been linked to resistance to doxorubicin^18^ and cisplatin^19^, suggesting it may contribute to stem-like characteristics. Despite these findings, a comprehensive understanding of the metastasis-related processes regulated by ELK3 in TNBC remains incomplete.

In this study, we employed lentiviral-mediated transduction to modulate ELK3 expression in TNBC cells, combined with whole transcriptome analysis to systematically map the global signaling networks influenced by ELK3. Functional analysis of ELK3-regulated gene sets enabled the prediction of metastasis-related cellular processes, which were subsequently validated through cellular assays in cells with either ELK3 knockdown or overexpression. Collectively, our findings identify ELK3 as a pro-metastatic transcription factor in TNBC that promotes cell motility and stemness.

## II. Materials and methods

### 1. Association between *ELK3* expression and metastasis-related traits of BC cell lines and tumors

RNA sequencing data for *ELK3* expression across 37 BC cell lines (CCLE_RNAseq_genes_rpkm_20180929.gct.gz) were retrieved from the Cancer Cell Line Encyclopedia (CCLE) (https://portals.broadinstitute.org/ccle). All 37 cell lines were further characterized for their EMT phenotype, by integrating the available data regarding their morphology in Matrigel^20^, in 2D^21^ and 3D^22^ models with the ATCC assignation (Suppl. Table 1). Differences in *ELK3* expression among phenotypes were analyzed using Kruskal-Wallis test, followed by Dunn’s multiple comparisons tests.

TCGA RNA sequencing data for *ELK3* (TCGA BRCA Dataset), along with corresponding clinicopathological data, were obtained from the Genomic Data Commons (https://portal.gdc.cancer.gov/). Differences in *ELK3* expression between clinical groups were evaluated using the Mann-Whitney U test.

### 2. Cell line and culture conditions

The TNBC cell line MDA-MB-231 (MDA231) was obtained from the European Collection of Authenticated Cell Cultures (ECACC) and maintained in RPMI-1640 medium supplemented with 10% fetal bovine serum (FBS), 1% penicillin–streptomycin, and 1% glutamine. All cell culture reagents were sourced from Gibco (Thermo Fisher Scientific).

### 3. Generation of genetically engineered cell lines by lentiviral-mediated transduction

MDA231-derived cell lines with either ELK3 overexpression (ELK3-OE) or knockdown (ELK3-KD), along with their respective mock controls (CTR-OE and CTR-KD), were generated using a third-generation lentiviral system, as previously described^23^. Briefly, plasmid cloning was performed in *Escherichia coli* DH5α, followed by viral packaging in HEK293T cells. Transfection of these cells was carried out for 24h with Lipofectamine 3000 (#L3000001, Thermo Fisher Scientific). Details of all plasmids used are provided in Supplementary Table 2. Viral infection of MDA231 cells was performed at 50% confluence, in the presence of 8 μg/μL polybrene (#TR-1003, Sigma-Aldrich). Transductions were performed in four biological replicates for each MDA231-derived cell line. Genetically modified cells were selected by culturing the cells in the presence of 4 μg/μL puromycin (#P8833, Sigma-Aldrich). Genetic transformation of the modified cell lines was validated by fluorescence microscopy (Observer D1 Inverted Fluorescence Microscope, Zeiss) and flow cytometry analysis (S3E Cell Sorter, Bio-Rad), through the detection of GFP protein.

### 4. Validation of the genetic modification by PCR

PCR analysis was conducted to detect the presence of the genetic construct or ELK3- specific sequences in the transformed cell lines. Genomic DNA was extracted using the PureLink Genomic DNA Kit (#K1820-01, Invitrogen). Target sequence amplification was performed on a MiniAmp ThermoCycler (A37834, Thermo Fisher Scientific) using the PCR MasterMix 2X kit (#K0171, Thermo Scientific), following the manufacturer’s instructions. Each PCR reaction was set up with 200 ng of DNA, a primer concentration of 0.5 μM, and a final reaction volume of 25 μL. The sequences, melting temperatures, and expected PCR product sizes for all primer pairs are provided in Supplementary Table 3.

### 5. Protein-level validation of the genetic modification by Western blot

Western blot analysis was performed to validate the altered ELK3 expression at the protein level in MDA231_ELK3-OE and MDA231_ELK3-KD cells. Briefly, a total of 20 μg of sample was loaded, proteins were separated on a 4–20% polyacrylamide gel (#4561094, Bio-Rad) for 1h and subsequently transferred onto a PVDF membrane (#1620177, Bio-Rad) for 2h. Primary antibodies used for detection included rabbit anti-Vinculin monoclonal antibody (#13901, Cell Signaling) and mouse anti-ELK3 monoclonal antibody (#MA5-25695, Invitrogen). Secondary antibodies used were Goat Anti-Rabbit IgG (#E-AB-1003, Elabscience) and Goat Anti-Mouse IgG (#E-AB-1001, Elabscience). All antibodies were applied at concentrations recommended by the manufacturers. Antigen-antibody complexes were visualized by chemiluminescence reaction (ChemiDoc Imaging System, BioRad). The protein expression was quantified by densitometry using ImageJ software. The statistical significance of differences in ELK3 protein expression between groups was assessed using t-test.

### 6. Evaluation of cell migration in 3D microfluidic devices

Microfluidic devices used for monitoring cell migration were fabricated by casting polydimethylsiloxane (PDMS, Dow Corning, Midland, MI) onto a microstructured mold created using standard photolithographic techniques, as previously described^24^. These devices consist of a central well connected to the surrounding medium through two arrays of 50 parallel channels, each measuring 600 µm in length and 10 µm in width. Both the central well and channels were coated with type IV collagen at a concentration of 20 µg/mL. A number of 4x10⁴ cells were seeded into the central well and the microfluidic device was placed on the stage of a microscope equipped with an incubation chamber (BioStation IMq, Nikon). Cell migration was monitored using time-lapse microscopy, capturing images every 10 minutes over a 24h period. Individual cell migration was analyzed using the Manual Tracking plugin in ImageJ software, based on two parameters: (1) migration speed, defined as the average displacement per 10-minute interval, and (2) migration velocity, calculated as the net displacement over the total distance traveled. The statistical significance of differences in migration capacity between groups was assessed using t-test.

### 7. Evaluation of cell cycle progression by flow-cytometry

Cell cycle progression was assessed by flow cytometry analysis of DNA content using DNA staining with propidium iodide (PI) (#V13242B, Thermo Fisher Scientific) for DNA content analysis. Briefly, cells were seeded in 6-well plates and allowed to adhere for 24h. Following incubation, cells were harvested by trypsinization, washed with cold PBS, and fixed in 70% ethanol. Subsequently, cells were permeabilized with 0.1% Triton X-100, and stained with PI (50 µg/mL) in a buffer containing 50 µg/mL RNase A (#46-7604, Invitrogen). Samples were analyzed using an S3E Cell Sorter (Bio-Rad) equipped with 488/561 nm lasers. The statistical significance of differences between groups was assessed using paired t-test.

### 8. Evaluation of cell proliferation by the AlamarBlue assay

The proliferation kinetics of all MDA231-derived cell lines was assessed over time using the AlamarBlue assay, following the manufacturer’s protocol. Briefly, 1x10⁴ cells/well were seeded in 200 µL of culture medium in 96-well plates, with four technical replicates per biological replicate. After a 2h incubation to allow cell adherence, 20 µL of AlamarBlue reagent (#DAL1025, Thermo Fisher Scientific) was added to each well. Absorbance was measured at 570 nm and 600 nm at 4, 8, 24, and 30h post-seeding. The statistical significance of differences between groups was assessed using paired t-test.

### 9. Evaluation of the cell stem-like phenotype by the mammosphere formation assay

Stemness potential of MDA231_ELK3-OE and MDA231_ELK3-KD cells was assessed using mammosphere formation assays. Cells were seeded at 1,000 cells per well in low-attachment 96-well plates, with four technical replicates per biological replicate. Each well contained 200 µL phenol-red free RPMI supplemented with glutamine (1%), penicillin (1%), non-essential amino acids (1%), N2 (1%), B27 (2%), EGF (10 ng/mL), and bFGF (20 ng/mL). Cultures were maintained for 7 days at 37°C under humidified conditions. Reagents were obtained from Gibco (Thermo Fisher Scientific). Mammospheres were visualized by phase-contrast microscopy using a Nikon AxioVision SE64 inverted microscope, and images were analyzed with ImageJ. Stemness was evaluated by measuring (1) average spheroid diameter and (2) the number of spheroids exceeding 50 µm and 60 µm per well. Statistical differences between groups were determined using a t-test.

### 10. Transcriptome analysis by microarray expression profiling

A whole transcriptome analysis using microarray technology was performed on all four biological replicates of MDA231_ELK3-KD and MDA231_CTR-KD cells. Cy-3-labeled microarray probes were synthesized from 50 ng of total RNA using the Low Input Quick Amp Labeling Kit (#5190-2331, Agilent), following the manufacturer’s protocol. Hybridization was carried out for 17h at 65°C on Agilent SurePrint G3 Human GE 8×60k arrays. Slides were scanned using an Agilent G2505C Microarray Scanner at 3 µm resolution, and microarray images were processed with Agilent Feature Extraction software v.11.5.1.1.

Preprocessing and differential expression analysis were conducted in R/Bioconductor using the raw median signals generated by FE. Control and flagged spots were removed, and data were quantile-normalized across arrays. A median signal value was computed for replicate probes on each array. Differential gene expression was assessed using linear models and empirical Bayes statistics, as implemented in the limma package^25^. Genes with ≥1.3-fold expression change and p<0.05 were considered significant. Data were deposited in the NCBI Gene Expression Omnibus (GEO) repository under accession number GSE302063.

Functional analyses were conducted using Ingenuity Pathway Analysis (IPA) and Gene Set Enrichment Analysis (GSEA). In IPA, the Disease and Function module predicted biological processes potentially activated or inhibited based on the activation z-score, with z-scores ≥2 or ≤-2 considered significant. In GSEA^26^, gene sets from both the Hallmark Gene Sets (h.all.v2024.1.Hs.symbols) and the Reactome Gene Sets (c2.cp.reactome.v2024.1.Hs.symbols) collections were considered enriched if they had a false discovery rate (FDR) below 25% and a nominal p-value < 0.01.

### 11. Gene expression analysis by RT-qPCR

Based on microarray data, the expression of randomly selected genes involved in cell migration, proliferation, and stemness was analyzed using RT-qPCR. Briefly, total RNA was extracted using the phenol-chloroform method. For each sample, 500 ng of RNA was reverse transcribed into cDNA using the RevertAid First Strand cDNA Synthesis Kit (#K1622, Thermo Scientific). PCR reactions were performed using the LightCycler TaqMan Master Kit (#04735536001, Roche) in a LightCycler 480 Thermocycler (Roche), following the manufacturer’s protocol. Primers and UPL probes used for amplification are detailed in Supplementary Table 3. Gene expression (fold change) was calculated using the ΔΔCt method^27^, with *RN18S1* serving as the housekeeping gene for normalization. Statistical significance between experimental groups was determined using t-test.

## III. Results

### 1. *ELK3* expression is associated with metastasis-related traits of BC cell lines and tumors

We previously quantified *ELK3* expression in seven BC cell lines (T47D, MCF7, MDA-MB-468, BT-549, HCC1937, HS578T, and MDA-MB-231) using RT-qPCR, and investigated its association with cell migration speed within 3D microfluidic devices^28^. A Spearman’s rank correlation analysis revealed a significant positive correlation between *ELK3* expression levels and the migratory capacity of these BC cell lines (Fig. 1A), suggesting that *ELK3* may serve as a key regulator of BC cell motility.

**Fig. 1.**
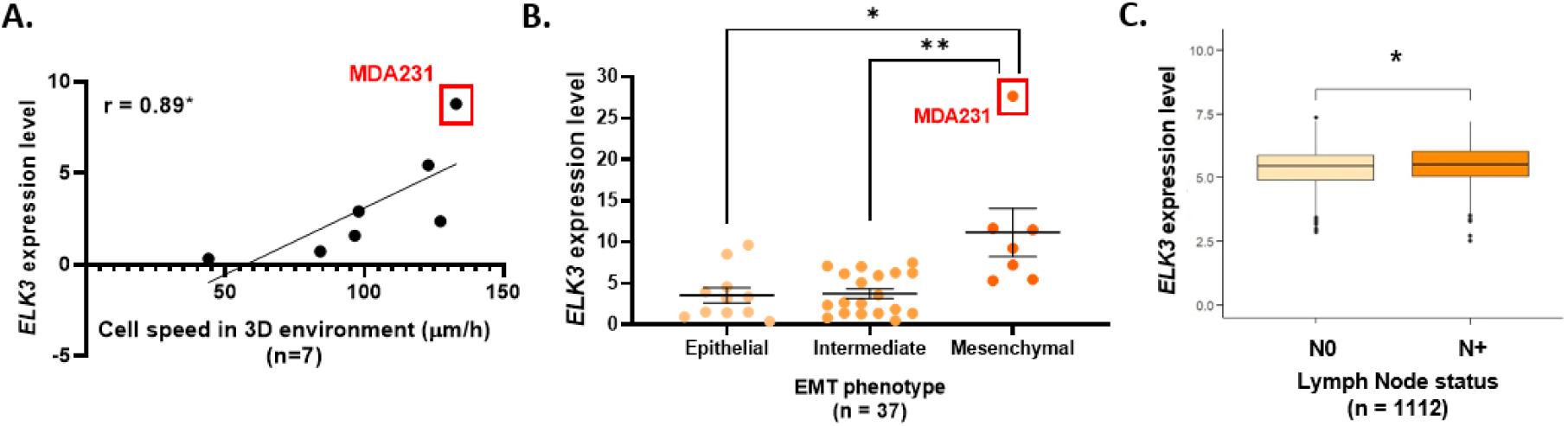
Association between *ELK3* expression and metastasis-related traits of BC cell lines and tumors: **A.** Spearman’s correlation between *ELK3* expression and the migration speed of seven BC cell lines in 3D environments; **B.** Association between *ELK3* expression (CCLE data) and EMT phenotype in 37 BC cell lines; **C.** Differential *ELK3* expression in breast cancer patients (TCGA data) stratified by lymph node status (N0: no lymph node invasion; N+: lymph node invasion). Statistical significance was assessed by Kruskal-Wallis test, followed by Dunn’s multiple comparisons tests (B) or Mann-Whitney test (C) (p*<0.05; p**<0.01; p***<0.001).

To further explore whether ELK3 expression contributes to the early stages of metastasis, its relationship with the EMT phenotype was examined in the present study across 37 BC cell lines (CCLE data). Cells with an epithelial phenotype exhibited predominantly low *ELK3* expression, whereas most mesenchymal cell lines displayed elevated *ELK3* levels. Statistically significant differences in *ELK3* expression were observed between epithelial/intermediate and mesenchymal phenotypes (Fig. 1B).

Clinical data further support the hypothesis that elevated *ELK3* expression is associated with increased metastatic potential. Analysis of a TCGA cohort comprising 1,112 BC patients, demonstrated significantly higher *ELK3* expression in tumors from patients with lymph node involvement compared to those without nodal metastasis (Fig. 1C)

### 2. Generation and characterization of genetically engineered TNBC cell lines with *ELK3* overexpression or knockdown

To investigate the role of ELK3 in metastasis-associated cellular processes, the MDA231 cell line was selected for both overexpression and knockdown. MDA231 is a mesenchymal cell line characterized by high migratory capacity, elevated *ELK3* expression (Fig. 1), and strong metastatic potential *in vivo*^29^.

Stable MDA231-derived cell lines with ELK3 overexpression (ELK3-OE) or knockdown (ELK3-KD), together with their respective mock controls (CTR-OE and CTR-KD), were generated using lentiviral transduction in four biological replicates. Following puromycin selection, more than 95% of cells exhibited successful genetic modification, as confirmed by GFP reporter expression (Fig. 2A, B). All transformed lines contained the vector constructs, while only MDA231_ELK3-OE and MDA231_ELK3-KD cells harbored the *ELK3* coding sequence or *ELK3* shRNA sequence, respectively (Fig. 2C). At the mRNA level, *ELK3* expression increased 1.78-fold in ELK3-OE cells and decreased 5.25-fold in ELK3-KD cells relative to their respective controls (Fig. 2D). Protein analysis confirmed these findings, showing a 2.29-fold increase in ELK3 protein in ELK3-OE cells and a 3.28-fold reduction in ELK3-KD cells (Fig. 2E). ELK3 transcriptional activity was assessed by measuring expression of the target gene *HIF1A*. Consistently, *HIF1A* expression was elevated in ELK3-OE cells and reduced in ELK3-KD cells (Fig. 2F).

**Fig. 2.**
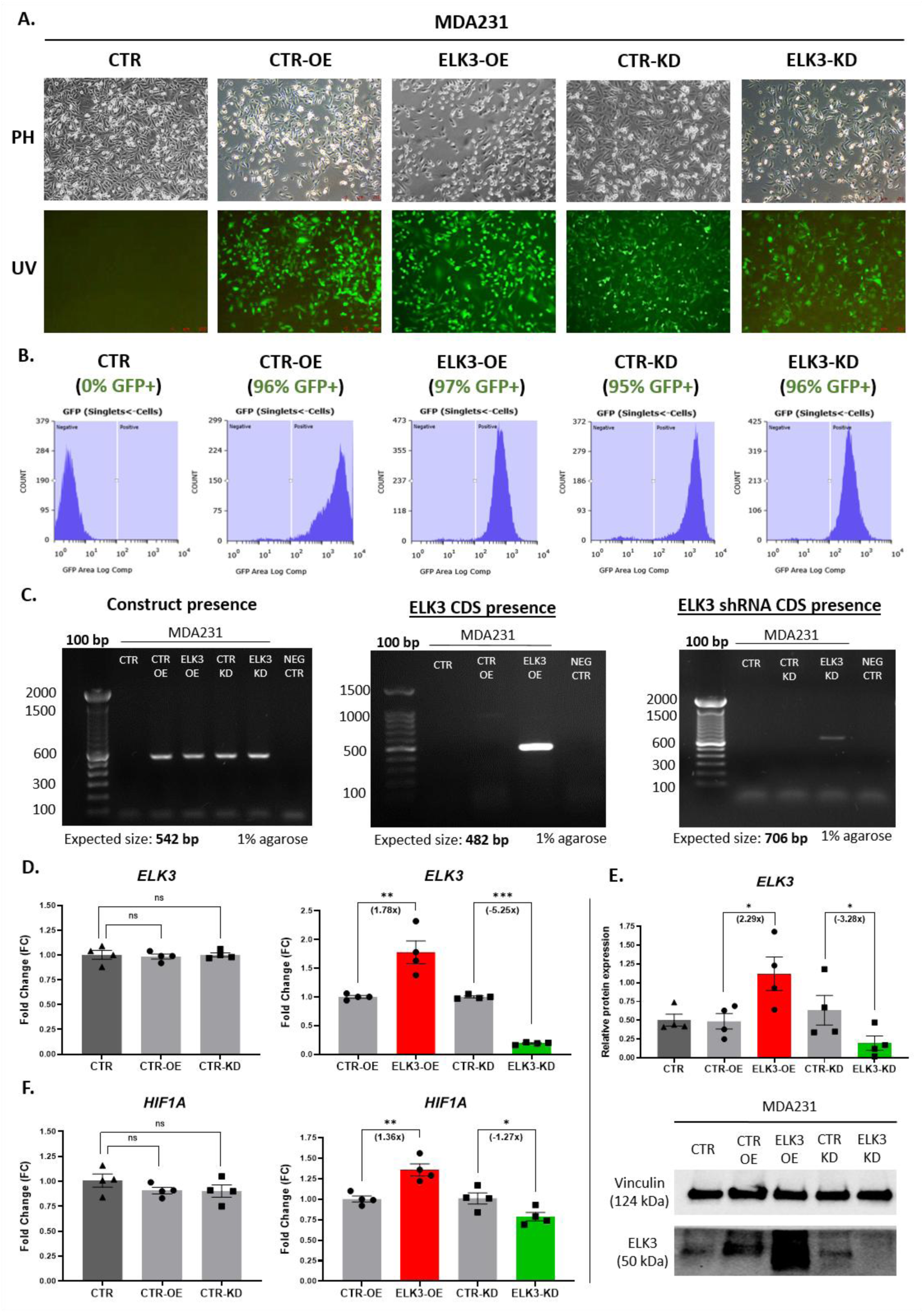
Characterization of genetically engineered MDA231-derived cell lines with ELK3 overexpression or knockdown: **A.** Fluorescence microscopy-based detection of *GFP* reporter gene expression to confirm genetic modification (representative biological replicate); **B.** Flow cytometry analysis of *GFP* expression to validate genetic modification (representative biological replicate); **C.** Genomic validation of genetic modification by PCR (representative biological replicate); **D.** Transcript-level confirmation of *ELK3* modulation using RT-qPCR (four biological replicates); **E.** Protein-level validation of ELK3 modulation by Western blot (four biological replicates); **F.** Expression levels of the ELK3 target gene *HIF1A* across engineered cell lines (four biological replicates). Statistical significance was assessed by t-test (p*<0.05; p**<0.01; p***<0.001).

To characterize molecular changes associated with *ELK3* loss-of-function, microarray-based whole transcriptome analysis was performed comparing ELK3-KD and CTR-KD cells (n=4). *ELK3* knockdown resulted in 2,156 significantly differentially expressed genes (p < 0.05), of which 740 exhibited a fold regulation greater than |±1.3| (Fig. 3A, B).

**Fig. 3.**
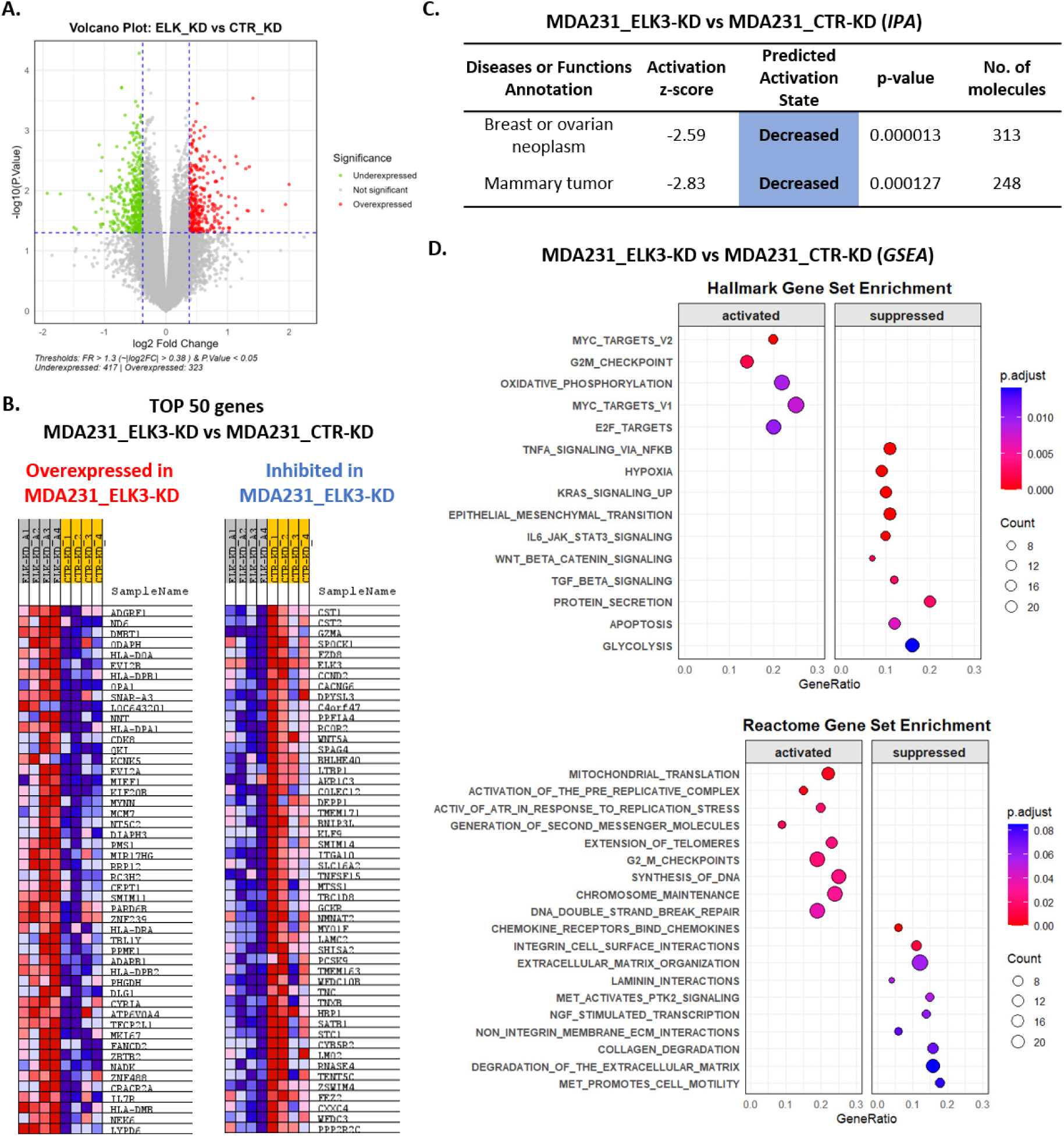
Summary of gene expression alterations following *ELK3* knockdown in MDA231 cells: **A.** Differentially expressed genes upon *ELK3* knockdown (FR>|±1.3|; p<0.05); **B.** Top 50 upregulated and downregulated genes after *ELK3* knockdown; **C.** IPA-based prediction of *ELK3* knockdown effects on breast cancer, based on microarray data; **D.** Main enriched cancer-related gene sets from Hallmark and Reactome Collections identified by GSEA, based on microarray data. The microarray experiments were performed on four biological replicates.

Pathway analysis (IPA) identified 43 significantly altered biological processes, including 26 cancer-related categories (Suppl. Table 4). Notably, the pathways associated with *Breast or ovarian neoplasm* and *Mammary tumor* were predicted to be significantly downregulated upon *ELK3* silencing (Fig. 3C). Additionally, *ELK3* inhibition impacted key cancer-related functional categories, including cellular movement, cell cycle regulation, cell death and survival (Suppl. Table 4). Consistent with these findings, GSEA revealed significant enrichment of 21 gene sets from the Hallmarks Collection and 104 from the Reactome Collection following *ELK3* knockdown (Suppl. Table 5). The top enriched cancer-related gene sets (Fig. 3D) were predominantly associated with processes such as cell migration and invasion, proliferation, apoptosis, and energy metabolism.

### 3. ELK3 enhances the migration of TNBC cells in 3D microfluidic systems

According to IPA analysis, *ELK3* silencing was predicted to inhibit seven cellular processes related to motility, including cell movement, adhesion and invasion of tumor cell lines (Fig. 4A; Suppl. Table 4). Additionally, processes involved in cell morphology, assembly, and structural organization, such as formation of focal adhesions and organization of actin cytoskeleton, both critical for migration, were also predicted to be suppressed (Fig. 4A).

**Fig. 4.**
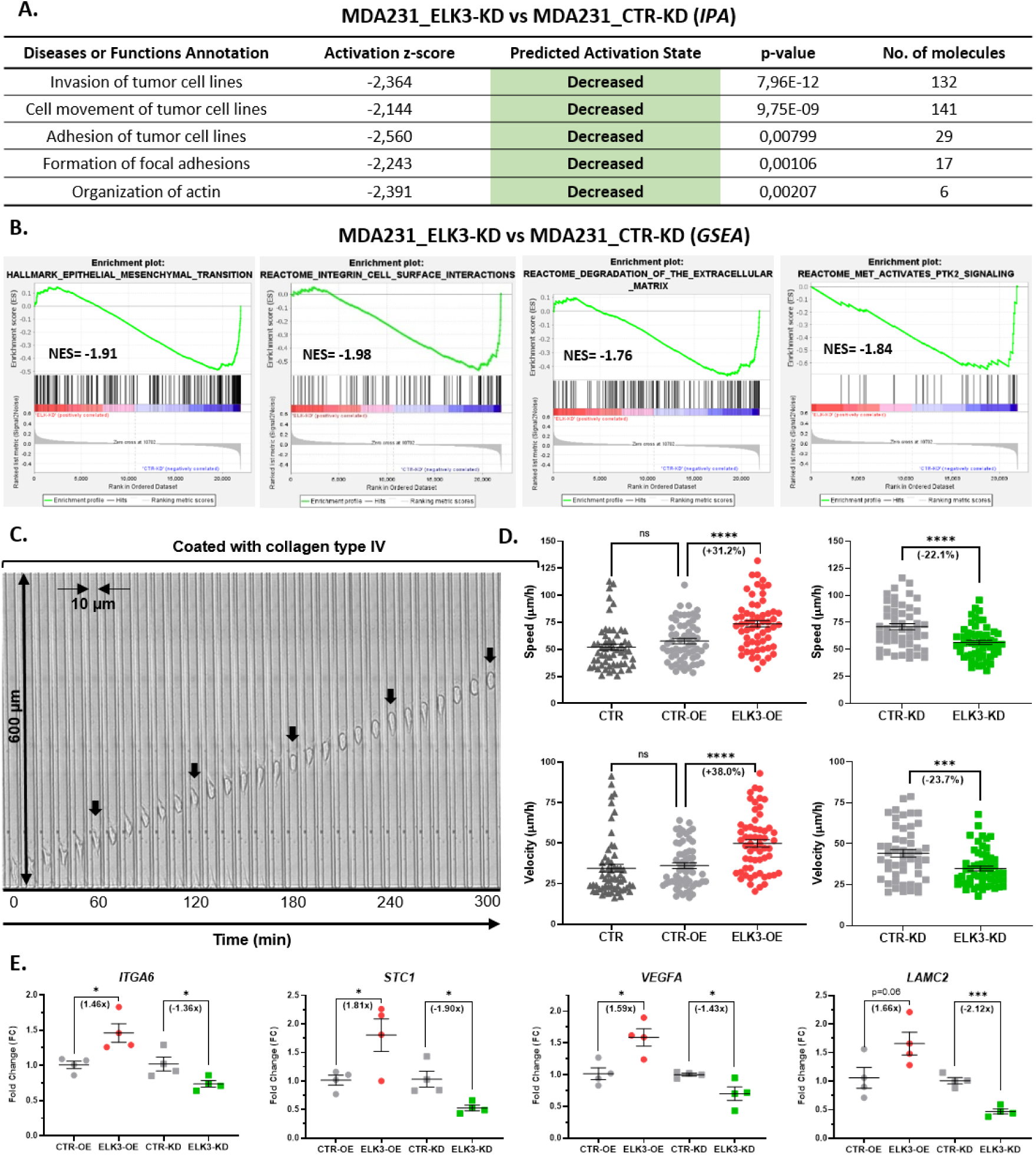
Impact of ELK3 on cell motility in MDA231 cells: **A.** Motility-associated cellular functions predicted to be affected by *ELK3* knockdown in IPA software, based on microarray data; **B.** Enriched motility-related gene sets from Hallmark and Reactome Collections following *ELK3* knockdown identified by GSEA, based on microarray data; **C.** Spatial characteristics of the microfluidic device used for the 3D cell migration assays; **D.** Migration capacity of parental MDA231 cells (CTR) and derived cell lines with *ELK3* overexpression (ELK3-OE) or knockdown (ELK3-KD), along with their respective mock controls (CTR-OE and CTR-KD), evaluated by migration speed and velocity in 3D microfluidic devices; **E.** Expression of selected motility-related genes in *ELK3*-overexpressing (ELK3-OE) and ELK3-knockdown (ELK3-KD) MDA231 cell lines, along with their respective mock controls (CTR-OE and CTR-KD), determined by RT-qPCR. All experiments were performed on four biological replicates. Statistical significance was assessed by t-test (p*<0.05; p**<0.01; p***<0.001).

Complementary results were obtained through GSEA, which showed that gene sets strongly associated with the promotion of cell migration and invasion, such as EMT, integrin-cell surface interactions and extracellular matrix (ECM) degradation, or signaling via the MET transcription factor, were enriched in MDA231_CTR-KD cells, indicating their downregulation upon *ELK3* knockdown (Fig. 4B).

In agreement with these molecular predictions, functional assays revealed a marked reduction in the migratory capacity of MDA231 cells following *ELK3* knockdown, as assessed in 3D microfluidic devices coated with collagen type IV (Fig. 4C). Migration speed and velocity decreased by approximately 22–23% after *ELK3* silencing (Fig. 4D). Conversely, MDA231 cells overexpressing *ELK3* demonstrated significantly enhanced migration, with a 31.2% increase in speed and a 38% increase in velocity (Fig. 4D).

To validate the microarray data, RT-qPCR analysis was performed on four key genes involved in cell motility: *ITGA6*, *STC1*, *VEGFA*, and *LAMC2*. Consistent with microarray results, *ELK3* knockdown suppressed the expression of all four genes (Fig. 4E). Notably, all genes were upregulated after *ELK3* overexpression (Fig. 4E).

### 4. ELK3 promotes a cancer stem-like phenotype in TNBC cells

IPA analysis predicted a reduction in the self-renewal capacity of MDA231 cells following *ELK3* knockdown, a hallmark feature of CSCs (Fig. 5A). Consistently, GSEA revealed that five key signaling pathways implicated in the regulation of cellular dedifferentiation and CSC formation, namely WNT/β-Catenin, JAK/STAT3, TGF-β, Notch1, and NF-κB, were enriched in MDA231_CTR-KD cells, suggesting their suppression upon *ELK3* silencing (Fig. 5B). Additionally, the hypoxia-associated gene set, representing a major driver of stemness acquisition, was significantly enriched in MDA231_CTR-KD cells, with a normalized enrichment score (NES) of -2.09 (Suppl. Table 5). The whole transcriptome data also suggested that angiogenesis, another driver of stemness, might be impaired following *ELK3* suppression, as indicated by the downregulation of multiple related processes, such as branching of endothelial cells (z***=-2.913) and formation of blood vessels (z**=-2.172) (Suppl. Table 4).

**Fig. 5.**
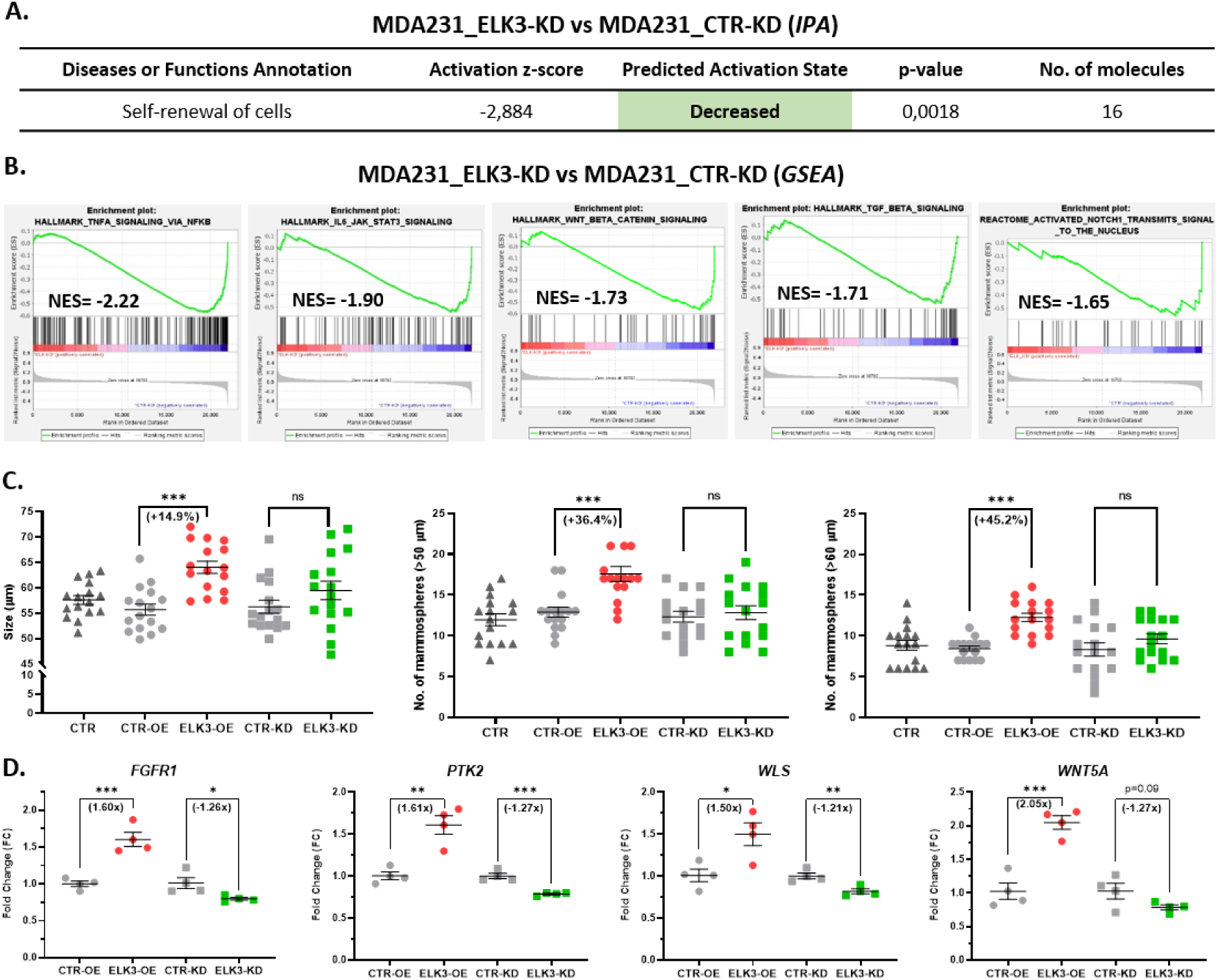
Impact of ELK3 on cancer cell stem-like phenotype in MDA231 cells: **A.** Stemness-associated cellular functions predicted to be affected by *ELK3* knockdown in IPA software, based on microarray data; **B.** Enriched stemness-related gene sets from Hallmark and Reactome Collections following *ELK3* knockdown identified by GSEA, based on microarray data; **C.** The stemness potential of parental MDA231 cells (CTR) and derived cell lines with *ELK3* overexpression (ELK3-OE) or knockdown (ELK3-KD), along with their respective mock controls (CTR-OE and CTR-KD), evaluated by the average mammosphere size and number in the mammosphere formation assay; **D.** Expression of selected stemness-related genes in *ELK3*-overexpressing (ELK3-OE) and ELK3-knockdown (ELK3-KD) MDA231 cell lines, along with their respective mock controls (CTR-OE and CTR-KD), determined by RT-qPCR. All experiments were performed on four biological replicates. Statistical significance was assessed by t-test (p*<0.05; p**<0.01; p***<0.001).

Supporting these molecular insights, *ELK3* overexpression in MDA231 cells led to a significant increase in stem-like potential, as shown by the mammosphere formation assay. Specifically, the average size of mammospheres increased by 14.9%, while the number of spheroids rose by 36.4% and 45.2% for structures exceeding 50 µm and 60 µm in diameter, respectively (Fig. 5C). Interestingly, *ELK3* silencing did not induce a significant decrease in mammosphere formation comparable to the enhancement seen with *ELK3* overexpression (Fig. 5C). This was unexpected given the transcriptomic data, suggesting the presence of compensatory mechanisms that may buffer the loss of *ELK3* in regulating stem-like traits.

To validate the microarray data, four genes implicated in CSC formation, namely *FGFR1*, *PTK2*, *WLS*, and *WNT5A*, were selected for further analysis. All four genes exhibited downregulation following *ELK3* knockdown (Fig. 5D). In contrast, their expression levels were significantly upregulated in MDA231 cells overexpressing *ELK3* (Fig. 5D).

### 5. ELK3 hampers cell proliferation in TNBC cells

IPA predicted an increase in processes related to cell proliferation, including cell cycle progression of tumor cell lines, following *ELK3* knockdown (Fig. 6A). In agreement, GSEA showed enrichment of signaling pathways associated with all phases of the cell cycle in MDA231_ELK3-KD cells, including the activation of the pre-replicative complex in G1, DNA synthesis during S phase, and progression through the G2/M checkpoint (Fig. 6B). Moreover, gene sets regulated by MYC and E2F, key drivers of the G1/S transition, were significantly enriched upon *ELK3* depletion (Fig. 6B). These results collectively suggest that ELK3 acts as a negative regulator of cell proliferation in MDA231 cells.

**Fig. 6.**
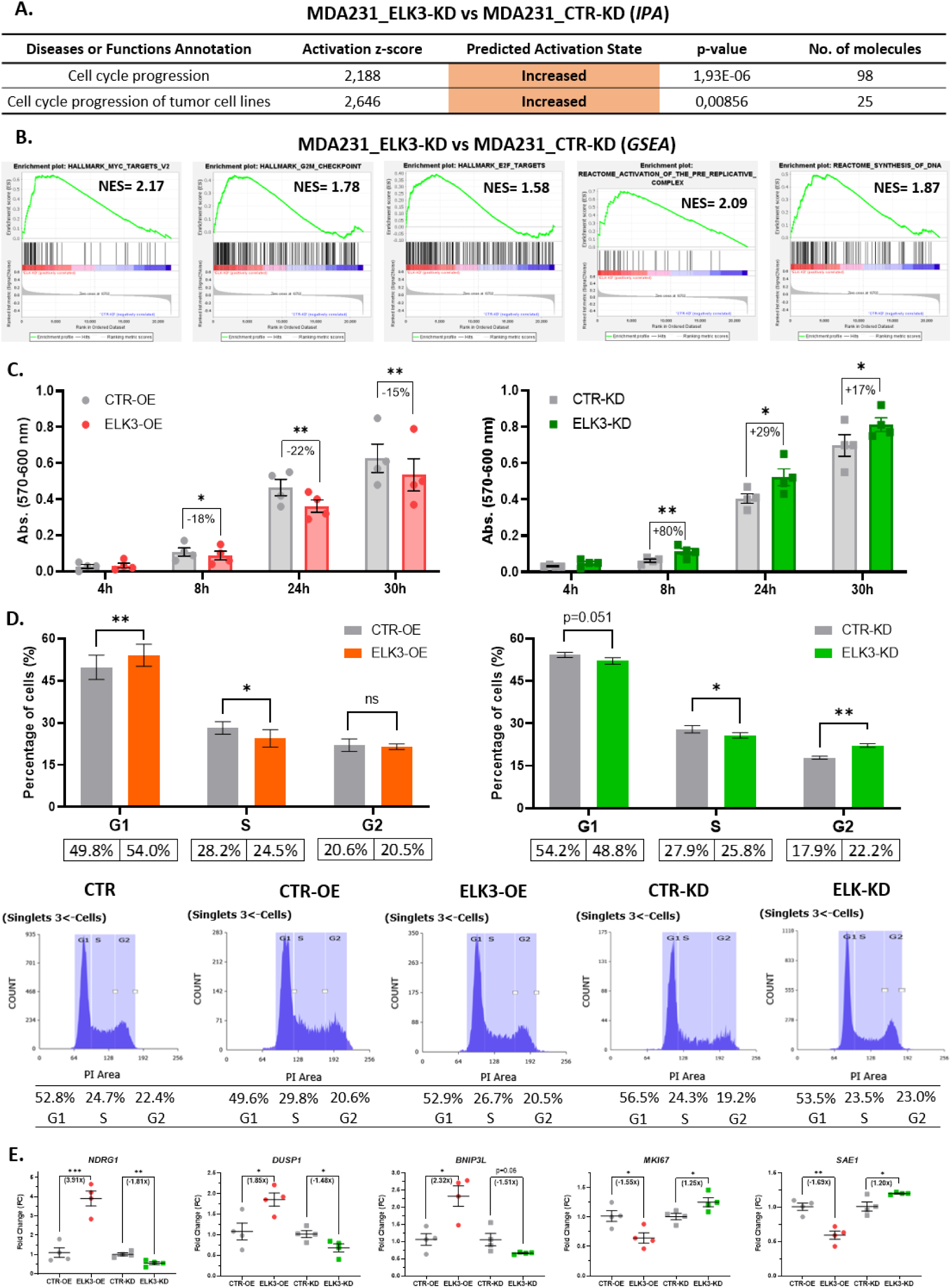
Impact of ELK3 on cell proliferation in MDA231 cells: **A.** Proliferation-associated cellular functions predicted to be affected by *ELK3* knockdown in IPA software, based on microarray data; **B.** Enriched proliferation-related gene sets from Hallmark and Reactome Collections following *ELK3* knockdown identified by GSEA, based on microarray data; **C.** Cell proliferation over time of *ELK3*-overexpressing (ELK3-OE) and *ELK3*-knockdown (ELK3-KD) MDA231 cell lines, in comparison with their respective mock controls (CTR-OE and CTR-KD), assessed by AlamarBlue assay; **D.** Cell cycle analysis in parental MDA231 cells (CTR) and derived cell lines with *ELK3* overexpression (ELK3-OE) or knockdown (ELK3-KD), along with their respective mock controls (CTR-OE and CTR-KD), evaluated by flow-cytometry; **E.** Expression of selected proliferation-related genes in *ELK3*-overexpressing (ELK3-OE) and ELK3-knockdown (ELK3-KD) MDA231 cell lines, along with their respective mock controls (CTR-OE and CTR-KD), determined by RT-qPCR. All experiments were performed on four biological replicates. Statistical significance was assessed by t-test (p*<0.05; p**<0.01; p***<0.001).

Consistent with the molecular findings, the AlamarBlue proliferation assay showed that *ELK3* knockdown resulted in a significant increase in cell number as early as 8h post-plating, while *ELK3* overexpression significantly suppressed cell proliferation starting at the same time point (Fig. 6C). In parallel, flow cytometry-based cell cycle analysis revealed that *ELK3* overexpression caused an accumulation of cells in the G1 phase, indicative of a G1/S transition block (Fig. 6D). Conversely, *ELK3* knockdown decreased the proportion of cells in G1 and significantly increased the G2 population (Fig. 6D). Taken together, these results support the conclusion that ELK3 impairs cell proliferation by arresting the cell cycle at the G1 phase.

Five genes associated with cell proliferation were selected to validate the microarray data by RT-qPCR. Genes known to hamper cell cycle progression, *NDRG1*, *DUSP1* and *BNIL3L*, were all downregulated following *ELK3* knockdown, whereas *MKI67* and *SAE1*, positive regulators of proliferation, were upregulated (Fig. 6E). In contrast, *ELK3* overexpression led to the inverse expression pattern for all five genes (Fig. 6E).

## IV. Discussion

Metastasis remains the leading cause of mortality in BC patients, yet no therapies specifically targeting the metastatic cascade are currently available in clinical practice. In TNBC, the most aggressive BC subtype, there are also no other approved targeted treatment options. Therefore, identifying novel molecular drivers of metastasis in TNBC is critical for advancing anti-metastatic therapeutic strategies and improving patient outcomes.

*ELK3* has previously been linked to TNBC, with significantly higher expression observed in basal-like and claudin-low BC cell lines compared to luminal and HER2-positive lines^10,14,15^. In terms of metastasis-associated traits, we have previously shown that *ELK3* expression strongly correlates with the migratory capacity of BC cells in 3D environments^28^. In this study, we further demonstrate that *ELK3* is associated with a mesenchymal phenotype, being significantly upregulated in mesenchymal-like BC cells relative to epithelial-like or intermediate phenotypes along the EMT spectrum. Moreover, analysis of patient samples revealed that *ELK3* expression was elevated in tumors from patients with lymph node involvement compared to those without nodal metastasis. Taken together, these findings support a role for ELK3 as a potential key driver of early metastatic events in TNBC.

Tumor cell migration and invasion are critical early events in the metastatic cascade, enabling a specific subset of cancer cells to detach from the primary tumor and infiltrate surrounding tissues. Our whole transcriptome analysis identified ELK3 as a regulator of cell motility in TNBC, predicting impaired migration and invasion upon *ELK3* knockdown. Consistent with these predictions, previous studies have shown that *ELK3* silencing reduces TNBC cell motility in 2D and endpoint assays^10,17^. Our results using 3D microfluidic systems, in both *ELK3* knockdown and overexpression models, provide further evidence of the pro-metastatic function of ELK3, by enhancing tumor cell migration in TNBC.

A range of additional processes relevant for cell migration and invasion were predicted to be regulated by ELK3, based on molecular profiling. GSEA analysis indicated that EMT, a key phenotypic switch enabling tumor cells to acquire migratory and invasive capabilities^30^, was downregulated in *ELK3* knockdown cells. Moreover, IPA results indicated that *ELK3* knockdown led to reduced cell adhesion, impaired focal adhesion formation, and disrupted actin cytoskeleton organization. In parallel, GSEA revealed that gene sets related to integrin-mediated cell surface interactions and ECM degradation were enriched in control cells. Interestingly, ELK3 also seemed to stimulate PTK2 (FAK) signaling, a pathway activated at integrin-mediated cell-ECM contact points that enhances both cell migration and stemness^31,32^. These findings suggest that ELK3 may promote EMT, enhance both cell–cell and cell–ECM adhesion, support the formation of cellular protrusions, and facilitate ECM degradation during tissue invasion. Consistent with these observations, previous studies on TNBC have shown that ELK3 inhibits E-cadherin expression^15^, inducing EMT^14^ and promotes cell motility by modulating cell–cell adhesion^16,17^, increasing filopodia-mediated migration through actin accumulation^33^, and indirectly upregulating MMP14 to drive ECM degradation^10^. Together, these findings potentially explain how ELK3 promotes tumor cell migration and invasion, underscoring its role as a key regulator of early metastatic events.

CSCs, a tumor subpopulation with self-renewal ability, play a crucial role in metastasis^13^. In addition to adopting a mesenchymal phenotype through EMT, metastatic cells often exhibit enhanced stem-like properties across various cancers, including breast cancer^34^. Although ELK3 signaling has not been previously associated with the stemness potential of TNBC cells, evidence from other cancer types suggests its involvement in regulating stem-like properties^11 12^. IPA functional analysis of our data indicated reduced self-renewal capacity in TNBC ELK3- silenced cells. Consistently, *ELK3* overexpression in MDA231 cells significantly increased stem-like potential, as demonstrated by clonogenic growth under non-adherent conditions.

CSC formation is driven by environmental factors such as hypoxia, abnormal angiogenesis, and chronic inflammation^13^. Our transcriptome analysis showed that hypoxia-related gene sets were significantly enriched in control MDA231_CTR-KD cells compared to *ELK3* knockdown cells, suggesting that ELK3 can mimic a hypoxic transcriptional response even under normoxic conditions. Additionally, *HIF1A* expression was upregulated in ELK3- overexpressing cells and downregulated following *ELK3* knockdown. Previous studies have shown that ELK3 regulates both the expression^35^ and stability^36^ of HIF1α, thereby contributing to the transcriptional hypoxia response^36^. Aberrant angiogenesis was also predicted to be reduced following *ELK3* suppression, with *VEGFA* expression increased with *ELK3* overexpression and decreased in *ELK3* knockdown cells. This aligns with previous findings linking ELK3 to angiogenesis, where the secretome of ELK3-KD TNBC cells was shown to inhibit peritumoral lymphangiogenesis *in vivo*^37^. Taken together, these findings suggest that ELK3 may enhance the stem-like properties of TNBC cells by inducing a hypoxia-like program and aberrant angiogenesis.

The primary intracellular pathways regulating stemness in cancer include the WNT/β- catenin, JAK/STAT3, TGF-β, Notch1, NF-κB, Hedgehog, PI3K/AKT and PPAR pathways^13^. Among these eight key signaling pathways, the GSEA analysis of our microarray data indicated that ELK3 activates the first five, highlighting the main molecular networks through which ELK3 may promote a stem-like phenotype in TNBC. Of these five ELK3-associated pathways, only NF-κB^37^ and TGF-β^14^ had previously been linked to ELK3 activity in breast cancer.

Another feature of CSCs is their quiescent state, which plays a critical role in conferring multidrug resistance. This quiescence, characterized by cell cycle arrest in the G0 phase, impedes proliferation and renders CSCs less susceptible to chemotherapeutic agents, which predominantly target actively dividing cells in the S and M phases^38^. Previous studies have demonstrated that ELK3 enhances chemoresistance to doxorubicin and cisplatin in TNBC^18,19^, suggesting a potential link between ELK3 activity and the quiescent state of the cells. In line with these findings, our data support the hypothesis that ELK3 promotes a CSC-like phenotype by also suppressing proliferation. The functional analysis of the whole transcriptome data in ELK3-KD cells predicted increased cell cycle activity in these cells, with significant enrichment of gene sets associated with the G1/S transition, including pathways regulated by MYC and E2F transcription factors and activation of the pre-replicative complex. Moreover, ELK3-KD cells exhibited enrichment of gene sets related to both the S phase (e.g., DNA synthesis) and M phase (e.g., G2/M checkpoint progression), indicating enhanced cell cycle dynamics. These computational predictions were corroborated by experimental evidence from cellular assays. Knockdown of ELK3 stimulated proliferation in TNBC cells, whereas overexpression of ELK3 suppressed it. The flow cytometry analysis further confirmed that ELK3-OE cells accumulated in the G0/G1 phase, reinforcing the role of ELK3 in inducing a quiescent state in the cells.

Previous studies investigating the role of ELK3 in regulating cell proliferation in BC have yielded seemingly contradictory results. In luminal BC cell lines, such as MCF7 and T47D, *ELK3* inhibition under adherent culture conditions led to a marked reduction in cell proliferation^39^. Conversely, in TNBC cells cultured under similar adherent conditions, *ELK3* knockdown was shown to enhance proliferation^14^, consistent with our findings. However, under non-adherent *in vitro* conditions or in *in vivo* models, *ELK3* knockdown in TNBC cells resulted in suppressed proliferation, manifested by smaller tumor sizes and reduced colony formation^14,35^. It is important to note that *in vivo* assays and *in vitro* assessments of anchorage-independent growth, while often interpreted through the lens of proliferation, primarily reflect the overall tumorigenic potential of cancer cells^40^, which is strongly associated with their stem-like properties^41^. Integrating these findings with our data, we propose that ELK3 suppresses proliferation under standard adherent culture conditions, but enhances the overall tumorigenicity of TNBC cells by promoting a CSC-like phenotype.

## V. Conclusion

ELK3 emerges as a key regulator of cellular processes implicated in early metastatic progression in TNBC. Elevated *ELK3* expression correlates with a mesenchymal phenotype in BC cell lines and with lymph node invasion in clinical tumor samples. Functionally, ELK3 enhances cell migration in 3D systems and drives the acquisition of CSC properties in TNBC. Within this CSC context, ELK3 supports both the self-renewal capacity and the quiescent state of cancer cells. These findings position ELK3 as a potential therapeutic target for disrupting the metastatic dissemination in TNBC.

