## Supplementary Table 1 for "Whole transcriptome analysis reveals ELK3 as a key driver of metastasis through regulation of 3D migration and stemness in triple-negative breast cancer cells"

**Supplementary Table 1**. The EMT phenotype of 37 breast cancer cell lines based on their morphology in Matrigel, in 2D and 3D models and the ATCC assignation

| **Cell line** | **Phenotype according to:** | | | | | **Phenotype (merged)** | **ELK3 expression** |
|---|---|---|---|---|---|---|---|
| | **Matrigel morphology [a]** | **3D culture [b]** | **2D culture [c]** | **cDNA microarray data [c]** | **ATCC [d]** | | |
| AU565_BREAST | grape | grape | | | | **intermediate** | 1.377 |
| BT474_BREAST | sphere | mass | epithelial | epithelial | | **epithelial** | 0.39904 |
| BT483_BREAST | sphere | mass | epithelial | epithelial | | **epithelial** | 3.16843 |
| BT549_BREAST | stellate | stellate | spindle | fibroblastic | mesenchymal-like | **mezenchymal** | 9.24976 |
| CAL120_BREAST | | | | | mesenchymal-like | **mezenchymal** | 11.49257 |
| CAL51_BREAST | | | | | mesenchymal-like | **mezenchymal** | 11.6609 |
| CAL851_BREAST | | | | | basal-like | **intermediate** | 7.47767 |
| CAMA1_BREAST | grape | grape | round | epithelial | | **intermediate** | 2.65615 |
| DU4475_BREAST | | | | epithelial | basal-like | **intermediate** | 5.9288 |
| HCC1143_BREAST | | | | | basal-like | **intermediate** | 7.02172 |
| HCC1187_BREAST | | | | | basal-like | **intermediate** | 5.0933 |
| HCC1500_BREAST | sphere | round | | | | **epithelial** | 3.93196 |
| HCC1569_BREAST | sphere | mass | | | | **epithelial** | 8.53249 |
| HCC1599_BREAST | | | | | basal-like | **intermediate** | 1.35089 |
| HCC1806_BREAST | | | | | basal-like | **intermediate** | 6.27626 |
| HCC1937_BREAST | | | epithelial | | basal-like | **intermediate** | 3.58374 |
| HCC2157_BREAST | | | | | basal-like | **intermediate** | 1.85307 |
| HCC38_BREAST | | | | | basal-like | **intermediate** | 6.16559 |
| HCC70_BREAST | sphere | mass | | | basal-like | **epithelial** | 9.63283 |
| HDQP1_BREAST | | | | | basal-like | **intermediate** | 6.25759 |
| HS578T_BREAST | stellate | stellate | spindle | fibroblastic | mesenchymal-like | **mezenchymal** | 7.21951 |
| MCF7_BREAST | sphere | mass | epithelial | epithelial | | **epithelial** | 1.51153 |
| MDAMB134VI_BREAST | grape | | round | epithelial | | **intermediate** | 1.3578 |
| MDAMB157_BREAST | | | spindle | | mesenchymal-like | **mezenchymal** | 5.47065 |
| MDAMB175VII_BREAST | sphere | | epithelial | epithelial | | **epithelial** | 4.59851 |
| MDAMB231_BREAST | stellate | stellate | spindle | fibroblastic | mesenchymal-like | **mezenchymal** | 27.65801 |

| | | | | | | | |
|---|---|---|---|---|---|---|---|
| MDAMB361_BREAST | sphere/grape | grape | epithelial | epithelial | | **intermediate** | 7.07626 |
| MDAMB415_BREAST | sphere | round | epithelial | | | **epithelial** | 1.50355 |
| MDAMB436_BREAST | stellate | stellate | spindle | | mesenchymal-like | **mezenchymal** | 5.3126 |
| MDAMB453_BREAST | grape | grape | round | epithelial | | **intermediate** | 1.29785 |
| MDAMB468_BREAST | sphere/grape | grape | round | | basal-like | **intermediate** | 2.55329 |
| SKBR3_BREAST | grape | grape | round | | | **intermediate** | 2.33238 |
| T47D_BREAST | sphere | mass | epithelial | epithelial | | **epithelial** | 1.43586 |
| UACC812_BREAST | grape | grape | epithelial | | | **intermediate** | 0.48673 |
| UACC893_BREAST | | | epithelial | | | **epithelial** | 3.35816 |
| ZR751_BREAST | sphere/grape | grape | epithelial | epithelial | | **epithelial** | 0.9452 |
| ZR7530_BREAST | | | round | | | **intermediate** | 0.82462 |

**The phenotype was determined by merging the descriptions of cell morphology, as presented in:**

[a] Blick T, Widodo E, Hugo H, Waltham M, Lenburg ME, Neve RM, Thompson EW. Epithelial mesenchymal transition traits in human breast cancer cell lines. Clin Exp Metastasis. 2008;25(6):629-42. doi: 10.1007/s10585-008-9170-6. Epub 2008 May 7. Review. PubMed PMID: 18461285.

[b] Kenny PA, Lee GY, Myers CA, Neve RM, Semeiks JR, Spellman PT, Lorenz K, Lee EH, Barcellos-Hoff MH, Petersen OW, Gray JW, Bissell MJ. The morphologies of breast cancer cell lines in three-dimensional assays correlate with their profiles of gene expression. Mol Oncol. 2007 Jun;1(1):84-96. doi: 10.1016/j.molonc.2007.02.004. PubMed PMID: 18516279; PubMed Central PMCID: PMC2391005.

[c] Hollestelle A, Peeters JK, Smid M, Timmermans M, Verhoog LC, Westenend PJ, Heine AA, Chan A, Sieuwerts AM, Wiemer EA, Klijn JG, van der Spek PJ, Foekens JA, Schutte M, den Bakker MA, Martens JW. Loss of E-cadherin is not a necessity for epithelial to mesenchymal transition in human breast cancer. Breast Cancer Res Treat. 2013 Feb;138(1):47-57. doi: 10.1007/s10549-013-2415-3. Epub 2013 Jan 22. PubMed PMID: 23338761.

[d] ATCC Breast Cancer Cell Panel (ATCC® 30-4500K)

The expression of *ELK3* gene in the breast cancer cel lines was obtained from CCLE
