## Supplementary Table 2 for "Whole transcriptome analysis reveals ELK3 as a key driver of metastasis through regulation of 3D migration and stemness in triple-negative breast cancer cells"

| Supplementary Table 2. Plasmids used for lentiviral-mediated trasnduction of MDA-MB-231 cells | | | | | |
|---|---|---|---|---|---|
| | **Type** | **Name** | **Product no.** | **Genes** | **Vector Size** |
| 1 | Packaging plasmid | pMDLg/pRRE | #12251, Addgene | Gag, Pol | 8890 bp |
| 2 | Regulatory plasmid | pRSV-Rev | #12253, Addgene | Rev | 4180 bp |
| 3 | Envelope plasmid | pMD2.G | #12259, Addgene | VSV-G | 5824 bp |
| 4 | Transfer plasmid - ELK3 overexpression | pLV[Exp]-EGFP:T2A:Puro-EF1A>hELK3 | #VB900001-4920cnr, VectorBuider | ELK3 CDS | 10598 bp |
| 5 | Transfer plasmid - ELK3 knock-down | pLV[shRNA]-EGFP:T2A:Puro-U6>hELK3[shRNA#1] | #VB900040-9805gbm, VectorBuider | ELK3-shRNA (Target sequence: ACTCGTCCTTCACCATTAATT) | 8347 bp |
| 6 | Transfer plasmid - mock control overexpression | pLV[Exp]-EGFP:T2A:Puro-EF1A>ORF_Stuffer | #VB010000-9389rbj, VectorBuider | ORF_Stuffer | 9623 bp |
| 7 | Transfer plasmid - mock control knock-down | pLV[shRNA]-EGFP:T2A:Puro-U6>Scramble_shRNA | #VB010000-0009mxc, VectorBuider | Scramble shRNA (Target sequence: CCTAAGGTTAAGTCGCCCTCG) | 8347 bp |
