## Supplementary Table 3 for "Whole transcriptome analysis reveals ELK3 as a key driver of metastasis through regulation of 3D migration and stemness in triple-negative breast cancer cells"

**Supplementary Table 3. Primers used in this study**

**2.1. Primers used for PCR for genome-level validation of the cell genetic modification**

| Name | Sequence (5'->3') | Tm | Expected PCR product size |
|---|---|---|---|
| **ELK-OE_F** | AAAACCGACAAGCACGTCAC | **59.6** | **482** |
| **ELK-OE_R** | AAGCAATCCATTTGGTGTCTGT | **58.7** | |
| **ELK-KD_F** | AAACACCGGACTCGTCCTTCA | **61.6** | **706** |
| **ELK-KD_R** | ACTTGTGGCCGTTTACGTCG | **61.2** | |
| **Const_F** | GTAAACGGCCACAAGTTCAGC | **60.3** | **542** |
| **Const_R** | CTCAGGTAGTGGTTGTCGGG | **59.8** | |

**2.2 Primers used for RT-qPCR**

| Name | Sequence (5'->3') | UPL |
|---|---|---|
| **ELK3_F** | AGCAGAGCCCTGCGATACTA | #29 |
| **ELK3_R** | TCTCCGGGAAGAGACAAAC | |
| **HIF1A_F** | GATAGCAAGACTTTCCTCAGTCG | #64 |
| **HIF1A_R** | TGGCTCATATCCCATCAATTC | |
| **ITGA6_F** | TTTGGCGTGGCTGACTTACA | #60 |
| **ITGA6_R** | GGAGGATGTCACCTGAGTGC | |
| **STC1_F** | TGCCCACCAAAGGTCAATGT | #52 |
| **STC1_R** | TAAGCGAGTTTGGAGTGGCC | |
| **VEGFA_F** | CTACCTCCACCATGCCAAGT | #29 |
| **VEGFA_R** | CCACTTCGTGATGATTCTGC | |
| **LAMC2_F** | AAGGGACTGGCCTCTCTGAA | #12 |
| **LAMC2_R** | TCGTGTCAAACTCCAGCTCC | |
| **FGFR1_F** | ACTCCGGCCTCTATGCTTG | #66 |
| **FGFR1_R** | AGGAGGGGAGAGCATCTGA | |
| **PTK2_F** | GAACTCTGACCTGGGTGAGC | #3 |
| **PTK2_R** | GTACTCTTGCTGGAGGCTGG | |
| **WLS_F** | ATCATCGCCTTTCTGGTGGG | #1 |
| **WLS_R** | ACCGACATGTAGGACACTGC | |
| **WNT5A_F** | ACGACCAGTTCAAGACCGTG | #59 |
| **WNT5A_R** | CTTGCACTTGACGTAGCAGC | |
| **NDRG1_F** | GGGTGCAGAAGGGACTAGG | #22 |
| **NDRG1_R** | TGCTCCTGGACATCAAACTCT | |
| **DUSP1_F** | CGAGGCCATTGACTTCATAGA | #65 |
| **DUSP1_R** | CTGGCAGTGGACAAACACC | |
| **BNIP3L_F** | AATGTCGTCCCACCTAGTCG | #70 |
| **BNIP3L_R** | AGCTCCACCCAGGAACTGT | |
| **MKI67_F** | GGACACGTGCCCAGAAAGTA | #17 |
| **MKI67_R** | GGTCTCCCCTGAGGTTTGTG | |
| **SAE1_F** | CTTGGCCCCAAGTGAACTCA | #45 |
| **SAE1_R** | GGAATACAGGTGGGCATGCT | |
