## Supplementary Table 5 for "Whole transcriptome analysis reveals ELK3 as a key driver of metastasis through regulation of 3D migration and stemness in triple-negative breast cancer cells"

| Supplementary Table 5.1 Hallmark gene sets enriched by ELK3 knockdown according to Gene Set Enrichment Analysis (GSEA) (NOM p-val<0.01; FDR<0.25) | | | | | | | |
|---|---|---|---|---|---|---|---|
| **Enriched in MDA231_ELK3-KD cell line** | | | | | | | |
| | **GS** | **SIZE** | **ES** | **NES** | **NOM p-val** | **FDR q-val** | **FWER p-val** |
| 1 | HALLMARK_MYC_TARGETS_V2 | 57 | 0.64 | 2.17 | 0 | 0 | 0 |
| 2 | HALLMARK_G2M_CHECKPOINT | 200 | 0.44 | 1.78 | 0 | 0.002 | 0.007 |
| 3 | HALLMARK_OXIDATIVE_PHOSPHORYLATION | 200 | 0.41 | 1.62 | 0 | 0.009 | 0.041 |
| 4 | HALLMARK_MYC_TARGETS_V1 | 199 | 0.4 | 1.62 | 0 | 0.008 | 0.044 |
| 5 | HALLMARK_E2F_TARGETS | 200 | 0.39 | 1.58 | 0 | 0.01 | 0.072 |
| **Enriched in MDA231_CTR-KD cell line** | | | | | | | |
| | **GS** | **SIZE** | **ES** | **NES** | **NOM p-val** | **FDR q-val** | **FWER p-val** |
| 1 | HALLMARK_TNFA_SIGNALING_VIA_NFKB | 199 | -0.57 | -2.22 | 0 | 0 | 0 |
| 2 | HALLMARK_HYPOXIA | 198 | -0.53 | -2.09 | 0 | 0 | 0 |
| 3 | HALLMARK_KRAS_SIGNALING_UP | 200 | -0.51 | -2 | 0 | 0 | 0 |
| 4 | HALLMARK_EPITHELIAL_MESENCHYMAL_TRANSITION | 200 | -0.49 | -1.91 | 0 | 0 | 0.001 |
| 5 | HALLMARK_IL6_JAK_STAT3_SIGNALING | 87 | -0.54 | -1.9 | 0 | 0 | 0.001 |
| 6 | HALLMARK_COMPLEMENT | 199 | -0.44 | -1.75 | 0 | 0.003 | 0.015 |
| 7 | HALLMARK_COAGULATION | 138 | -0.46 | -1.73 | 0 | 0.003 | 0.015 |
| 8 | HALLMARK_WNT_BETA_CATENIN_SIGNALING | 42 | -0.57 | -1.73 | 0 | 0.002 | 0.015 |
| 9 | HALLMARK_UV_RESPONSE_DN | 143 | -0.45 | -1.72 | 0 | 0.002 | 0.017 |
| 10 | HALLMARK_TGF_BETA_SIGNALING | 54 | -0.53 | -1.71 | 0.004 | 0.003 | 0.021 |
| 11 | HALLMARK_PROTEIN_SECRETION | 96 | -0.47 | -1.67 | 0.002 | 0.003 | 0.03 |
| 12 | HALLMARK_APOPTOSIS | 161 | -0.43 | -1.6 | 0.002 | 0.007 | 0.07 |
| 13 | HALLMARK_INFLAMMATORY_RESPONSE | 200 | -0.41 | -1.59 | 0 | 0.007 | 0.078 |
| 14 | HALLMARK_GLYCOLYSIS | 198 | -0.39 | -1.52 | 0.002 | 0.014 | 0.155 |
| 15 | HALLMARK_MYOGENESIS | 198 | -0.38 | -1.5 | 0.003 | 0.017 | 0.191 |
| 16 | HALLMARK_FATTY_ACID_METABOLISM | 156 | -0.39 | -1.46 | 0.004 | 0.023 | 0.269 |

| Supplementary Table 5.2 Reactome gene sets enriched by ELK3 knockdown according to Gene Set Enrichment Analysis (GSEA) (NOM p-val<0.01; FDR<0.25) | | | | | | | |
|---|---|---|---|---|---|---|---|
| **Enriched in MDA231_ELK3-KD cell line** | | | | | | | |
| | **GS** | **Size** | **ES** | **NES** | **NOM p-val** | **FDR q-val** | **FWER p-val** |
| 1 | REACTOME_MITOCHONDRIAL_TRANSLATION | 94 | 0.58 | 2.11 | 0 | 0.004 | 0.004 |
| 2 | REACTOME_ACTIVATION_OF_THE_PRE_REPLICATIVE_COMPLEX | 33 | 0.7 | 2.09 | 0 | 0.003 | 0.005 |
| 3 | REACTOME_RRNA_MODIFICATION_IN_THE_NUCLEUS_AND_CYTOSOL | 60 | 0.62 | 2.08 | 0 | 0.002 | 0.005 |
| 4 | REACTOME_PD_1_SIGNALING | 22 | 0.77 | 2.07 | 0 | 0.002 | 0.008 |

| | | | | | | | |
|---|---|---|---|---|---|---|---|
| 5 | REACTOME_RRNA_PROCESSING | 201 | 0.48 | 1.95 | 0 | 0.018 | 0.079 |
| 6 | REACTOME_TRNA_PROCESSING | 107 | 0.52 | 1.92 | 0 | 0.023 | 0.115 |
| 7 | REACTOME_ACTIVATION_OF_ATR_IN_RESPONSE_TO_REPLICATION_STRESS | 37 | 0.62 | 1.92 | 0 | 0.02 | 0.115 |
| 8 | REACTOME_GENERATION_OF_SECOND_MESSENGER_MOLECULES | 33 | 0.65 | 1.91 | 0 | 0.018 | 0.122 |
| 9 | REACTOME_RNA_POLYMERASE_III_TRANSCRIPTION_INITIATION_FROM_TYPE_1_PROMOTER | 28 | 0.66 | 1.91 | 0 | 0.018 | 0.13 |
| 10 | REACTOME_EXTENSION_OF_TELOMERES | 51 | 0.57 | 1.88 | 0 | 0.023 | 0.183 |
| 11 | REACTOME_G2_M_CHECKPOINTS | 163 | 0.48 | 1.88 | 0 | 0.022 | 0.196 |
| 12 | REACTOME_SYNTHESIS_OF_DNA | 121 | 0.5 | 1.87 | 0 | 0.024 | 0.237 |
| 13 | REACTOME_TELOMERE_MAINTENANCE | 107 | 0.51 | 1.86 | 0 | 0.023 | 0.246 |
| 14 | REACTOME_CHROMOSOME_MAINTENANCE | 134 | 0.48 | 1.83 | 0 | 0.029 | 0.379 |
| 15 | REACTOME_SNRNP_ASSEMBLY | 53 | 0.55 | 1.81 | 0 | 0.034 | 0.45 |
| 16 | REACTOME_DNA_DOUBLE_STRAND_BREAK_REPAIR | 164 | 0.45 | 1.81 | 0 | 0.037 | 0.497 |
| 17 | REACTOME_DNA_REPLICATION | 181 | 0.45 | 1.8 | 0 | 0.037 | 0.504 |
| 18 | REACTOME_COSTIMULATION_BY_THE_CD28_FAMILY | 68 | 0.52 | 1.8 | 0 | 0.035 | 0.508 |
| 19 | REACTOME_ORC1_REMOVAL_FROM_CHROMATIN | 71 | 0.52 | 1.8 | 0 | 0.034 | 0.511 |
| 20 | REACTOME_TRANSLATION | 292 | 0.43 | 1.8 | 0 | 0.035 | 0.544 |
| 21 | REACTOME_DNA_REPAIR | 324 | 0.41 | 1.75 | 0 | 0.054 | 0.722 |
| 22 | REACTOME_S_PHASE | 163 | 0.45 | 1.74 | 0 | 0.053 | 0.771 |
| 23 | REACTOME_DNA_REPLICATION_PRE_INITIATION | 153 | 0.44 | 1.71 | 0 | 0.069 | 0.886 |
| 24 | REACTOME_PROCESSING_OF_CAPPED_INTRON_CONTAINING_PRE_MRNA | 281 | 0.41 | 1.71 | 0 | 0.068 | 0.891 |
| 25 | REACTOME_SWITCHING_OF_ORIGINS_TO_A_POST_REPLICATIVE_STATE | 92 | 0.48 | 1.7 | 0 | 0.067 | 0.899 |
| 26 | REACTOME_MITOTIC_G1_PHASE_AND_G1_S_TRANSITION | 149 | 0.43 | 1.69 | 0 | 0.069 | 0.92 |
| 27 | REACTOME_MHC_CLASS_II_ANTIGEN_PRESENTATION | 122 | 0.45 | 1.69 | 0 | 0.069 | 0.928 |
| 28 | REACTOME_CELL_CYCLE_CHECKPOINTS | 287 | 0.41 | 1.69 | 0 | 0.068 | 0.932 |
| 29 | REACTOME_HOMOLOGY_DIRECTED_REPAIR | 134 | 0.44 | 1.68 | 0 | 0.068 | 0.946 |
| 30 | REACTOME_SUMOYLATION_OF_UBIQUITINYLATION_PROTEINS | 38 | 0.55 | 1.66 | 0 | 0.079 | 0.976 |
| 31 | REACTOME_TRNA_PROCESSING_IN_THE_NUCLEUS | 57 | 0.5 | 1.66 | 0 | 0.078 | 0.977 |
| 32 | REACTOME_CITRIC_ACID_CYCLE_TCA_CYCLE | 34 | 0.55 | 1.65 | 0 | 0.082 | 0.985 |
| 33 | REACTOME_RESPIRATORY_ELECTRON_TRANSPORT | 144 | 0.42 | 1.64 | 0 | 0.088 | 0.992 |
| 34 | REACTOME_HCMV_LATE_EVENTS | 108 | 0.43 | 1.6 | 0 | 0.109 | 0.999 |
| 35 | REACTOME_MRNA_SPLICING | 212 | 0.39 | 1.57 | 0 | 0.119 | 1 |

| | | | | | | | |
|---|---|---|---|---|---|---|---|
| 36 | REACTOME_PROTEIN_LOCALIZATION | 163 | 0.39 | 1.56 | 0 | 0.124 | 1 |
| 37 | REACTOME_MEIOSIS | 113 | 0.41 | 1.53 | 0 | 0.139 | 1 |
| 38 | REACTOME_M_PHASE | 407 | 0.34 | 1.47 | 0 | 0.172 | 1 |
| 39 | REACTOME_DNA_STRAND_ELONGATION | 32 | 0.62 | 1.88 | 0.002 | 0.021 | 0.199 |
| 40 | REACTOME_RECYCLING_OF_BILE_ACIDS_AND_SALTS | 18 | 0.71 | 1.86 | 0.002 | 0.024 | 0.277 |
| 41 | REACTOME_MITOCHONDRIAL_PROTEIN_IMPORT | 65 | 0.54 | 1.84 | 0.002 | 0.026 | 0.332 |
| 42 | REACTOME_G0_AND_EARLY_G1 | 27 | 0.61 | 1.74 | 0.002 | 0.052 | 0.757 |
| 43 | REACTOME_G1_S_SPECIFIC_TRANSCRIPTION | 29 | 0.59 | 1.72 | 0.002 | 0.062 | 0.839 |
| 44 | REACTOME_INTERACTIONS_OF_REV_WITH_HOST_CELLULAR_PROTEINS | 36 | 0.54 | 1.68 | 0.002 | 0.07 | 0.945 |
| 45 | REACTOME_TRANSCRIPTIONAL_REGULATION_BY_SMALL_RNAS | 99 | 0.44 | 1.61 | 0.002 | 0.1 | 0.999 |
| 46 | REACTOME_BASE_EXCISION_REPAIR | 85 | 0.45 | 1.59 | 0.002 | 0.108 | 1 |
| 47 | REACTOME_G2_M_DNA_DAMAGE_CHECKPOINT | 90 | 0.44 | 1.56 | 0.002 | 0.124 | 1 |
| 48 | REACTOME_DNA_DAMAGE_TELOMERE_STRESS_INDUCED_SENESCENCE | 74 | 0.44 | 1.55 | 0.002 | 0.128 | 1 |
| 49 | REACTOME_HIV_INFECTION | 229 | 0.34 | 1.37 | 0.002 | 0.239 | 1 |
| 50 | REACTOME_NUCLEAR_PORE_COMPLEX_NPC_DISASSEMBLY | 35 | 0.61 | 1.86 | 0.004 | 0.023 | 0.283 |
| 51 | REACTOME_CRISTAE_FORMATION | 31 | 0.62 | 1.8 | 0.004 | 0.034 | 0.521 |
| 52 | REACTOME_TRNA_MODIFICATION_IN_THE_NUCLEUS_AND_CYTOSOL | 43 | 0.55 | 1.76 | 0.004 | 0.053 | 0.697 |
| 53 | REACTOME_NUCLEAR_IMPORT_OF_REV_PROTEIN | 33 | 0.6 | 1.75 | 0.004 | 0.054 | 0.736 |
| 54 | REACTOME_TELOMERE_C_STRAND_LAGGING_STRAND_SYNTHESIS | 34 | 0.58 | 1.74 | 0.004 | 0.054 | 0.748 |
| 55 | REACTOME_RNA_POLYMERASE_III_CHAIN_ELONGATION | 18 | 0.66 | 1.74 | 0.004 | 0.053 | 0.755 |
| 56 | REACTOME_EXPORT_OF_VIRAL_RIBONUCLEOPROTEINS_FROM_NUCLEUS | 32 | 0.56 | 1.69 | 0.004 | 0.069 | 0.923 |
| 57 | REACTOME_DUAL_INCISION_IN_TC_NER | 62 | 0.48 | 1.64 | 0.004 | 0.085 | 0.989 |
| 58 | REACTOME_DNA_DOUBLE_STRAND_BREAK_RESPONSE | 74 | 0.47 | 1.66 | 0.005 | 0.079 | 0.974 |
| 59 | REACTOME_PROCESSING_OF_DNA_DOUBLE_STRAND_BREAK_ENDS | 93 | 0.45 | 1.62 | 0.005 | 0.093 | 0.996 |
| 60 | REACTOME_TRANSCRIPTION_COUPLED_NUCLEOTIDE_EXCISION_REPAIR_TC_NER | 75 | 0.47 | 1.62 | 0.005 | 0.093 | 0.996 |
| 61 | REACTOME_TRANSPORT_OF_MATURE_TRANSCRIPT_TO_CYTOPLASM | 82 | 0.44 | 1.56 | 0.005 | 0.123 | 1 |
| 62 | REACTOME_HCMV_EARLY_EVENTS | 128 | 0.38 | 1.45 | 0.005 | 0.19 | 1 |
| 63 | REACTOME_HCMV_INFECTION | 152 | 0.37 | 1.43 | 0.005 | 0.197 | 1 |
| 64 | REACTOME_MITOTIC_METAPHASE_AND_ANAPHASE | 234 | 0.34 | 1.4 | 0.005 | 0.221 | 1 |

| | | | | | | | |
|---|---|---|---|---|---|---|---|
| 65 | REACTOME_NONHOMOLOGOUS_END_JOINING_NHEJ | 64 | 0.45 | 1.54 | 0.007 | 0.133 | 1 |
| 66 | REACTOME_MITOCHONDRIAL_BIOGENESIS | 94 | 0.42 | 1.5 | 0.007 | 0.156 | 1 |
| 67 | REACTOME_HIV_LIFE_CYCLE | 147 | 0.38 | 1.45 | 0.007 | 0.189 | 1 |
| 68 | REACTOME_AEROBIC_RESPIRATION_AND_RESPIRATORY_ELECTRON_TRANSPORT | 244 | 0.35 | 1.42 | 0.007 | 0.206 | 1 |
| **Enriched in MDA231_CTR-KD cell line** | | | | | | | |
| | **GS** | **SIZE** | **ES** | **NES** | **NOM p-val** | **FDR q-val** | **FWER p-val** |
| 1 | REACTOME_CHEMOKINE_RECEPTORS_BIND_CHEMOKINES | 56 | -0.68 | -2.2 | 0 | 0 | 0 |
| 2 | REACTOME_REGULATION_OF_INSULIN_LIKE_GROWTH_FACTOR_IGF_TRANSPORT_AND_UPTAKE_BY_INSULIN_LIKE_GROWTH_FACTOR_BINDING_PROTEINS_IGFBPS | 122 | -0.58 | -2.15 | 0 | 0 | 0 |
| 3 | REACTOME_INTEGRIN_CELL_SURFACE_INTERACTIONS | 85 | -0.57 | -1.98 | 0 | 0.01 | 0.03 |
| 4 | REACTOME_COMMON_PATHWAY_OF_FIBRIN_CLOT_FORMATION | 22 | -0.7 | -1.87 | 0.002 | 0.061 | 0.24 |
| 5 | REACTOME_EXTRACELLULAR_MATRIX_ORGANIZATION | 299 | -0.46 | -1.86 | 0 | 0.057 | 0.267 |
| 6 | REACTOME_LAMININ_INTERACTIONS | 30 | -0.64 | -1.85 | 0 | 0.053 | 0.294 |
| 7 | REACTOME_MET_ACTIVATES_PTK2_SIGNALING | 30 | -0.65 | -1.84 | 0.002 | 0.053 | 0.334 |
| 8 | REACTOME_NGF_STIMULATED_TRANSCRIPTION | 39 | -0.61 | -1.82 | 0 | 0.062 | 0.417 |
| 9 | REACTOME_NON_INTEGRIN_MEMBRANE_ECM_INTERACTIONS | 58 | -0.55 | -1.79 | 0 | 0.075 | 0.559 |
| 10 | REACTOME_COLLAGEN_DEGRADATION | 64 | -0.53 | -1.79 | 0 | 0.072 | 0.579 |
| 11 | REACTOME_DOWNREGULATION_OF_TGF_BETA_RECEPTOR_SIGNALING | 26 | -0.64 | -1.79 | 0.004 | 0.077 | 0.536 |
| 12 | REACTOME_TGF_BETA_RECEPTOR_SIGNALING_ACTIVATES_SMADS | 47 | -0.57 | -1.77 | 0.004 | 0.083 | 0.659 |
| 13 | REACTOME_DEGRADATION_OF_THE_EXTRACELLULAR_MATRIX | 140 | -0.47 | -1.76 | 0 | 0.085 | 0.693 |
| 14 | REACTOME_NUCLEAR_EVENTS_KINASE_AND_TRANSCRIPTION_FACTOR_ACTIVATION | 61 | -0.54 | -1.76 | 0.002 | 0.083 | 0.715 |
| 15 | REACTOME_MET_PROMOTES_CELL_MOTILITY | 41 | -0.57 | -1.75 | 0.002 | 0.083 | 0.736 |
| 16 | REACTOME_ASSEMBLY_OF_COLLAGEN_FIBRILS_AND_OTHER_MULTIMERIC_STRUCTURES | 61 | -0.52 | -1.74 | 0 | 0.083 | 0.778 |
| 17 | REACTOME_ACYL_CHAIN_REMODELLING_OF_PC | 26 | -0.62 | -1.74 | 0.005 | 0.082 | 0.751 |
| 18 | REACTOME_ADHERENS_JUNCTIONS_INTERACTIONS | 57 | -0.53 | -1.72 | 0 | 0.093 | 0.838 |
| 19 | REACTOME_INTERLEUKIN_10_SIGNALING | 45 | -0.54 | -1.72 | 0 | 0.092 | 0.861 |
| 20 | REACTOME_MATURATION_OF_SPIKE_PROTEIN | 36 | -0.57 | -1.72 | 0.006 | 0.094 | 0.853 |
| 21 | REACTOME_ACYL_CHAIN_REMODELLING_OF_PI | 17 | -0.67 | -1.7 | 0.006 | 0.111 | 0.918 |

| | | | | | | | |
|---|---|---|---|---|---|---|---|
| 22 | REACTOME_FORMATION_OF_FIBRIN_CLOT_CLOTTING_CASCADE | 39 | -0.56 | -1.69 | 0.002 | 0.113 | 0.933 |
| 23 | REACTOME_SIGNALING_BY_MET | 79 | -0.49 | -1.66 | 0.002 | 0.133 | 0.984 |
| 24 | REACTOME_ACTIVATED_NOTCH1_TRANSMITS_SIGNAL_TO_THE_NUCLEUS | 31 | -0.56 | -1.65 | 0.008 | 0.123 | 0.992 |
| 25 | REACTOME_ACYL_CHAIN_REMODELLING_OF_PS | 21 | -0.64 | -1.65 | 0.009 | 0.129 | 0.989 |
| 26 | REACTOME_ACYL_CHAIN_REMODELLING_OF_PE | 28 | -0.59 | -1.65 | 0.009 | 0.125 | 0.991 |
| 27 | REACTOME_RAC1_GTPASE_CYCLE | 184 | -0.42 | -1.63 | 0 | 0.134 | 0.995 |
| 28 | REACTOME_ARACHIDONIC_ACID_METABOLISM | 59 | -0.5 | -1.63 | 0.007 | 0.137 | 0.995 |
| 29 | REACTOME_CELL_JUNCTION_ORGANIZATION | 116 | -0.44 | -1.61 | 0 | 0.146 | 0.997 |
| 30 | REACTOME_COLLAGEN_FORMATION | 90 | -0.46 | -1.61 | 0.005 | 0.143 | 0.997 |
| 31 | REACTOME_INTERLEUKIN_4_AND_INTERLEUKIN_13_SIGNALING | 108 | -0.45 | -1.6 | 0.004 | 0.144 | 0.998 |
| 32 | REACTOME_RAC3_GTPASE_CYCLE | 94 | -0.45 | -1.58 | 0.005 | 0.163 | 1 |
| 33 | REACTOME_LATE_SARS_COV_2_INFECTION_EVENTS | 69 | -0.47 | -1.57 | 0.005 | 0.168 | 1 |
| 34 | REACTOME_ECM_PROTEOGLYCANS | 76 | -0.45 | -1.55 | 0.007 | 0.19 | 1 |
| 35 | REACTOME_CELL_CELL_JUNCTION_ORGANIZATION | 89 | -0.43 | -1.53 | 0.007 | 0.195 | 1 |
| 36 | REACTOME_CELL_CELL_COMMUNICATION | 154 | -0.4 | -1.51 | 0 | 0.214 | 1 |
