## Supplementary Table 4 for "Whole transcriptome analysis reveals ELK3 as a key driver of metastasis through regulation of 3D migration and stemness in triple-negative breast cancer cells"

**Supplementary Table 4.** Statistically significant processes modulated by ELK3 knockdown according to Ingenuity Pathway Analysis software - Disease and Function Annotation (z > +/- 2)

© 2000-2024 QIAGEN. All rights reserved.

| Diseases or Functions Annotation | p-value | Predicted Activation State | Activation z-score | # Molecules |
|---|---|---|---|---|
| Breast or ovarian neoplasm | 0.0000138 | Decreased | -2.59 | 313 |
| Mammary tumor | 0.000127 | Decreased | -2.83 | 248 |
| Shape change of vascular endothelial cells | 0.000985 | Decreased | -2.963 | 9 |
| Branching of vasculature | 0.00455 | Decreased | -2.5 | 11 |
| Branching of endothelial cells | 0.000105 | Decreased | -2.913 | 12 |
| Sprouting of vascular endothelial cells | 0.000274 | Decreased | -2.792 | 8 |
| Formation of blood vessel | 0.0063 | Decreased | -2.172 | 16 |
| Senescence of cells | 0.000031 | Decreased | -2.684 | 38 |
| Cell cycle progression | 0.00000193 | Increased | 2.188 | 98 |
| Cell cycle progression of tumor cell lines | 0.00856 | Increased | 2.646 | 25 |
| Cell survival | 0.000248 | Decreased | -2.133 | 133 |
| Cell viability | 0.000278 | Decreased | -2.098 | 128 |
| Cell viability of breast cancer cell lines | 0.000628 | Decreased | -2.117 | 26 |
| Cell viability of tumor cell lines | 0.000712 | Decreased | -2.203 | 89 |
| Self-renewal of cells | 0.0018 | Decreased | -2.884 | 16 |
| Apoptosis of neurons | 0.00861 | Increased | 2.018 | 37 |
| Apoptosis of sarcoma cell lines | 0.00895 | Increased | 2.088 | 18 |
| Apoptosis of epithelial cells | 0.0114 | Increased | 2.532 | 27 |
| Sprouting | 0.0034 | Decreased | -2.524 | 42 |
| Branching of cells | 0.00223 | Decreased | -2.733 | 41 |
| Adhesion of tumor cell lines | 0.00799 | Decreased | -2.56 | 29 |
| Binding of tumor cell lines | 0.0117 | Decreased | -2.459 | 35 |
| Interaction of tumor cell lines | 0.0118 | Decreased | -2.575 | 36 |
| Formation of focal adhesions | 0.00106 | Decreased | -2.243 | 17 |
| Phagocytosis of cells | 0.00237 | Decreased | -2.148 | 32 |
| Organization of actin | 0.00207 | Decreased | -2.391 | 6 |
| Invasion of tumor cell lines | 7.96E-12 | Decreased | -2.364 | 132 |
| Invasion of cells | 6.13E-11 | Decreased | -2.282 | 143 |
| Cell movement of tumor cell lines | 9.75E-09 | Decreased | -2.144 | 141 |
| Cell movement of sarcoma cell lines | 0.000495 | Decreased | -3.127 | 17 |
| Migration of sarcoma cell lines | 0.00075 | Decreased | -2.49 | 15 |
| Cell movement of melanoma cell lines | 0.00973 | Decreased | -2.184 | 15 |
| Cell movement of bone cancer cell lines | 0.0113 | Decreased | -2.437 | 10 |
| Growth Failure | 0.000831 | Increased | 2.566 | 52 |
| Growth failure or short stature | 0.00301 | Increased | 2.648 | 59 |
| Development of head | 0.00436 | Decreased | -2.034 | 86 |
| Development of genitourinary system | 0.00502 | Decreased | -2.086 | 80 |
| Immune response of cells | 0.000832 | Decreased | -2.138 | 53 |
| Motor dysfunction or movement disorder | 0.00361 | Increased | 2.343 | 104 |
| Development of exocrine gland | 0.00399 | Decreased | -2.543 | 21 |
| Organismal death | 8.25E-08 | Increased | 3.384 | 227 |
| Respiratory failure | 0.00753 | Increased | 2.225 | 15 |
| Branching of epithelial tissue | 0.0023 | Decreased | -2.913 | 13 |
